# Distinct neural mechanisms meet challenges in dynamic visual attention due to either load or object spacing

**DOI:** 10.1101/472209

**Authors:** V. Mäki-Marttunen, T. Hagen, B. Laeng, T. Espeseth

## Abstract

When solving dynamic visuo-spatial tasks, the brain copes with perceptual and cognitive processing challenges. In the multiple-object tracking (MOT) task, the number of objects to be tracked (i.e. load) imposes attentional demands, but so does spatial interference from irrelevant objects (i.e. crowding). Presently, it is not clear whether load and crowding activate separate cognitive and physiological mechanisms. Such knowledge would be important to understand the neurophysiology of visual attention. Furthermore, it would help resolve conflicting views between theories of visual cognition, particularly concerning sources of capacity limitations. To address this problem, we varied the degree of processing challenge in the MOT task in two ways: First, the number of objects to track, and second, the spatial proximity between targets and distractors. We first measured task-induced pupil dilations and saccades during MOT. In a separate cohort we measured fMRI brain activity during MOT. The behavioral results in both cohorts revealed that increased load and crowding led to reduced accuracy in an additive manner. Load was associated with pupil dilations, whereas crowding was not. Activity in dorsal attentional areas and frequency of saccades were proportionally larger both with higher levels of load and crowding. Higher crowding recruited additionally ventral attentional areas that may reflect orienting mechanisms. The activity in the brainstem nuclei ventral tegmental area/substantia nigra and locus coeruleus showed clearly dissociated patterns. Our results constitute convergent evidence from independent samples that processing challenges due to load and object spacing may rely on different mechanisms.

## 1. Introduction

When faced with a visuo-spatial task that requires top-down control -that is, a task that cannot be solved solely automatically-, the logistics of attention become engaged. The brain continuously assesses the appropriate level of processing resources and recruits optimally the mechanisms that deal with current demands. In the case of multiple-object tracking (MOT, Z. W. Pylyshyn and Storm (1988)), the brain flexibly handles the attentional challenge posed by the number of moving objects to be tracked simultaneously (“load”; Meyerhoff et al. (2017); Scimeca and Franconeri (2015)). Subsequently, the brain copes with emerging situations of spatial interference during the tracking period. However, whether the same or different brain mechanisms are involved in dealing with both types of challenges has not yet been established.

Different accounts were formulated to explain how attention deals with the different challenges during multiple-object tracking. In addition to the number of targets, other factors influence accuracy: the speed of the objects (Feria, 2013; Meyerhoff et al., 2016; Tombu & Seiffert, 2008), the duration of tracking (Oksama & Hyönä, 2004), the distance the objects travel (S. Franconeri et al., 2010), and the frequency of close encounters between targets and distractors (a concept known as “crowding”, (Alvarez & Franconeri, 2007; 2012; Feria, 2013; S. L. Franconeri et al., 2008; He et al., 1996; Iordanescu et al., 2009; Shim et al., 2008)). All these factors seem to be associated with the degree of proximity between targets and distractors. One theoretical account proposes that the number of close encounters (i.e. events with reduced inter-object spacing) is indeed the only limiting factor in MOT (S. Franconeri et al., 2010; Scimeca & Franconeri, 2015), and the limitation to attention may relate to the ability of keeping differentiated mental representations of the target objects (Drew et al., 2013; Drew et al., 2012; Drew & Vogel, 2008). A different view would be that both load and crowding rely on distinct processes. In this line, recent studies propose that the brain is able to deal with close encounters by flexibly distributing attentional resources (Iordanescu et al., 2009; Meyerhoff et al., 2016; Srivastava & Vul, 2016). The underlying system can achieve this by either altering the spatial distribution of attention covertly or by directing the eyes in order to gain higher spatial resolution in areas undergoing crowding (e.g. rescue saccades, Zelinsky and Todor (2010)). This perspective proposes that load may modulate the amount of available resources, and re-allocations of attentional resources increase as the minimum distance between objects gets smaller. The flexible resource is supposed to be dynamically allocated to increase the local attentional resolution for targets requiring higher precision processing (Iordanescu, Grabowecky, & Suzuki, 2009; Meyerhoff, Papenmeier, Jahn, & Huff, 2016; Srivastava & Vul, 2016). Thus, an increase in number of targets engages increasing resources, and object spacing should become more critical for performance with increased number of targets. The limitation to attention may be, in addition to keeping a mental representation of the targets, the ability to flexibly allocate resources (for example, attention or working memory, Allen et al. (2006); Lapierre et al. (2017); Vul et al. (2009)).

To look closer at the two possibilities accounting for limitations in MOT, we designed a MOT task where both load and crowding were manipulated so that any unique or interactive effects may be scrutinized. Specifically, in addition to the load manipulation (2, 3 and 4 targets), we varied the amount of crowding within the trials, with trials of low or high crowding. Importantly, in the high crowding condition the number of close encounters between targets and distractors was equalized across load levels; that is, there was the same number of close encounters at each load level. In order to evaluate whether specific physiological responses are associated with each type of challenge, we leveraged ocular measures and brain activity in two separate experiments with independent cohorts. In the eye-tracking experiment we measured the pupil size and the frequency of saccades during tracking. The brainstem locus coerules-norepinephrine (LC-NE) system has been strongly tied to pupil dilations, effort, and arousal, and has been suggested to be involved in an attentional filter that selects for the temporal occurrence of relevant objects (Aston-Jones & Cohen, 2005; Joshi et al., 2016). In MOT, the LC-NE system is recruited with higher load (Alnæs et al., 2014; Wright et al., 2013). A still open question is whether instances of close encounters during tracking also activate this system. Spatial attention has been associated with the dopaminergic system (DA, Noudoost and Moore (2011); Thiele and Bellgrove (2018)), produced in the brainstem nuclei ventral tegmental area and substantia nigra (VTA/SN). For example, direct infusion of dopaminergic agents in the frontal eye fields (FEF) influence saccades and visual processing in macaques (Noudoost & Moore, 2011), and dopamine has potent modulatory effects on prefrontal activation to spatial working memory tasks (Arnsten (1997); Sawaguchi and Goldman-Rakic (1991)). In addition, the DA system has been implicated in effort, (Husain & Roiser, 2018; Pessiglione et al., 2018; Westbrook & Braver, 2016; Westbrook et al., 2015), but may be more strongly associated with valuation and effort-based decision-making than active attentional engagement during task performance (Varazzani et al., 2015). What is more, inter-individual variability in performance in MOT (Drew & Vogel, 2008; Oksama & Hyönä, 2004; Störmer et al., 2013) may be subject to differences in the recruitment of those effort-related systems (Unsworth & Robison, 2015). Assessing the activity in the LC-NE and VTA/SN-DA systems allowed us to evaluate the dynamic allocation of attention during MOT due to increasing load and crowding.

Recent neuroimaging research has revealed that a fronto-parietal brain network is recruited during the tracking of multiple objects, showing increased activity with increasing load (Alnæs et al., 2014; Culham et al., 1998; Culham et al., 2001; Jahn et al., 2012; Jovicich et al., 2001; Shim et al., 2009; Tomasi et al., 2004). This is consistent with findings that implicate this network in dealing with attentional demands across tasks and modalities (Corbetta & Shulman, 2002; Duncan, 2010; Ptak, 2012). It would be expected that both the number of objects to track and the events of crowding should engage an effort-related system, but it remains unclear what is the specificity of this system.

In summary, we pursued the question of whether load and crowding constitute the same or distinct attentional challenges. Furthermore, we asked how the neuromodulatory and attentional systems of the brain instantiate the attentional mechanisms by which the brain deals with load and crowding. Given that crowding underlies load effects (S. Franconeri et al., 2010), a critical prediction was that there should be no effect of the load manipulation when crowding is kept constant (Figure 1.a). Manipulating either load or crowding would recruit the same cortical and subcortical regions. Furthermore, we expected crowding to increase the frequency of saccades and the activity of the VTA/SN. Importantly, we expected that crowding would also trigger pupil dilations and activity in the LC, but there should be no effect of load on these measures when crowding is kept constant. Alternatively, if load and crowding impose at least partially distinct limitations to the brain, we would expect performance to drop with load at constant crowding levels (Figure 1.b-c), and a distinction at the physiological and neural level. Pupil dilation and LC activity would increase as a function of load, even when crowding is kept constant. Moreover, while a general demand network would be engaged parametrically with load to sustain and update the representation of the objects, additional activation may help dealing with arising events of close encounters. This would imply a more fine-regulated processing that provides more specificity and efficacy. Finally, we tested whether individual differences in performance can be associated to distinct patterns of recruitment of the neuromodulatory systems.

**Figure 1.**
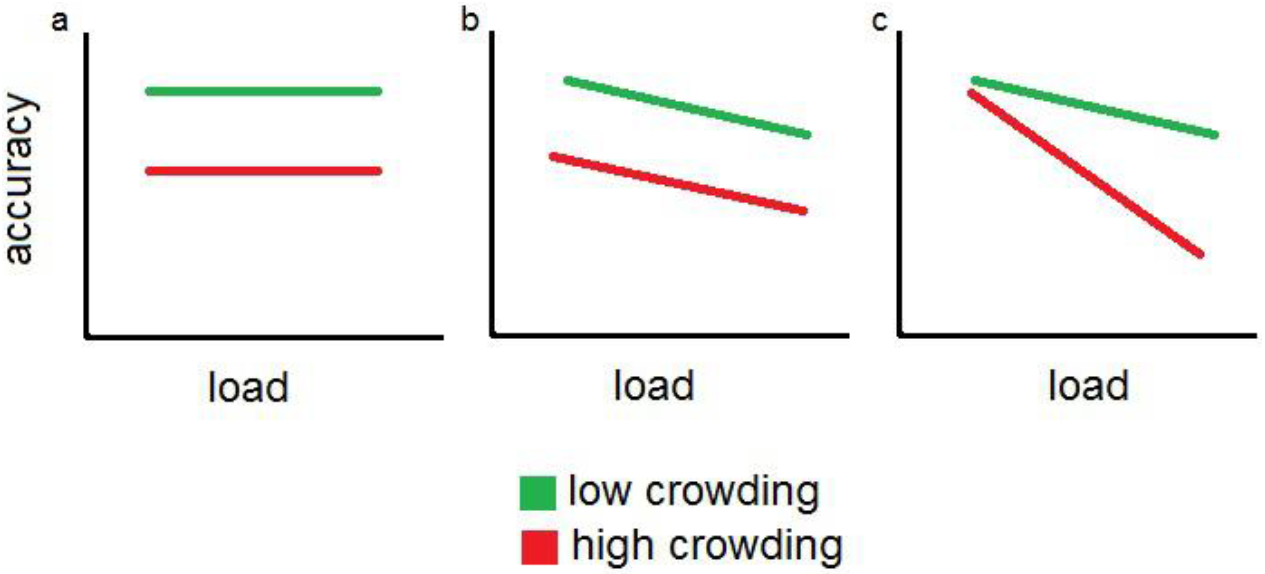
Hypothesized effects of load and crowding on accuracy. If in each crowding level (low/high) the amount of crowding (i.e. close encounters) is held constant across load levels, the hypothesized patterns differ whether load and crowding rely on the same or different mechanisms: a) expected pattern for the load effect being given by the amount of crowding; b) expected pattern for a load-crowding model with independent mechanisms; c) expected pattern for a load-crowding model with shared mechanisms. This would be expected because the two loading factors would act as “dual-tasks” (Strobach et al., 2018; Williges & Wierwille, 1979).

## 2. Materials and Methods

### 2.1 MOT task

In the MOT task (Figure 2), participants are presented with 10 objects (disks). Some of them are cued as targets by changing their color for a short period. Afterwards, these targets acquire the same appearance as the other objects (or distractors) and then all the disks start moving in random directions, changing paths only when bouncing with each other or the borders of the display area. The participant’s task is to track the targets as the objects move around and, at the end of the trial, indicate which objects in the display belonged to the target set. We parametrically varied the number of target objects, or cognitive load, so that the participants should correspondingly adjust the required degree of attentional effort. In the present experiments, we used three levels of load, with trials of two, three or four objects to be tracked. In addition, we included two levels of crowding for each of the load levels: low and high crowding. Crowding is defined as the number of instances where one distractor approaches a target within a certain close-range distance (see description below). This proximity increases confusion about the target, therefore increasing the risk of losing the target or swapping the target with a distractor.

**Figure 2.**
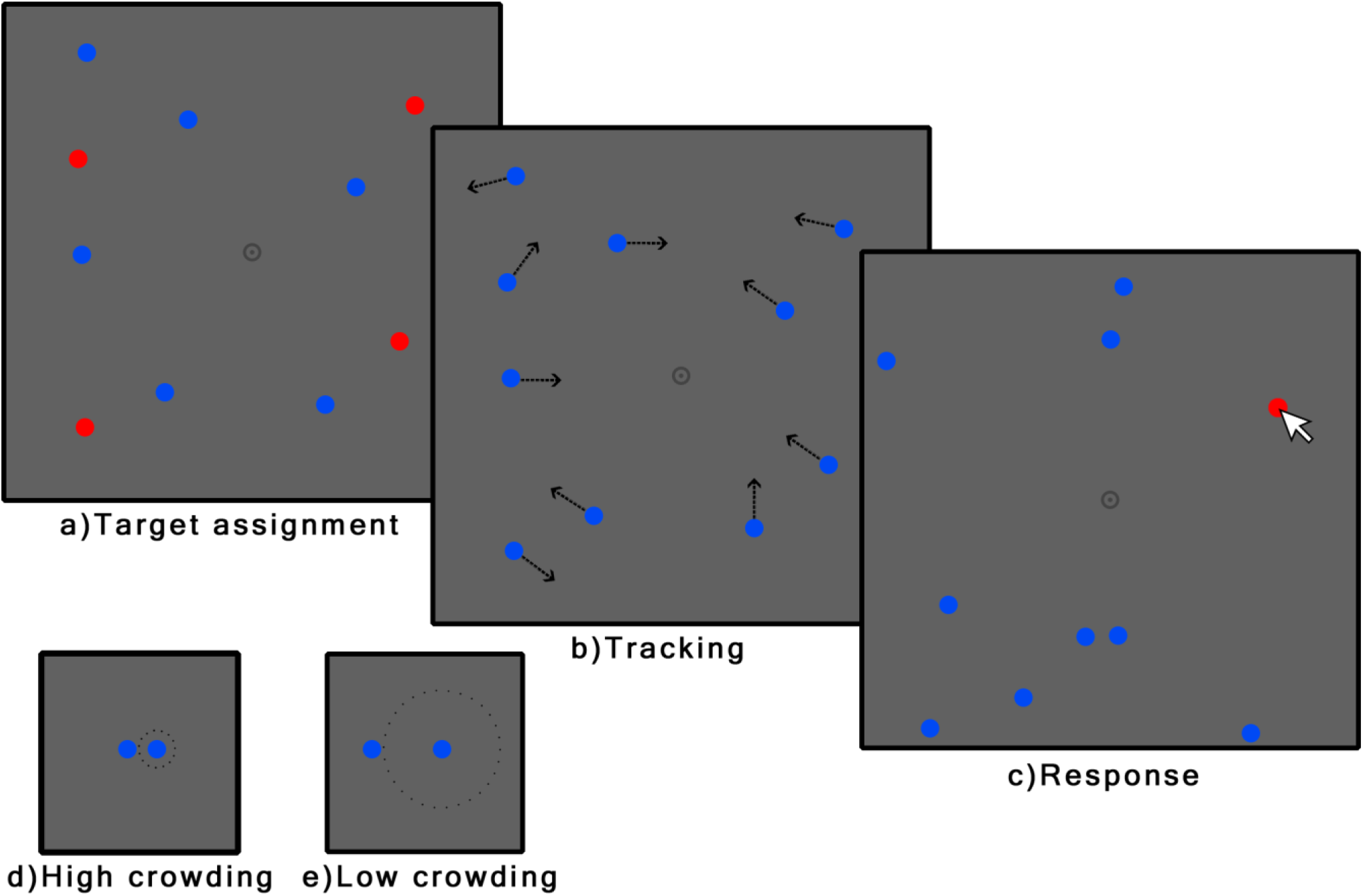
MOT task. In the assignment period of a trial (a), ten black disks are presented. Then, a number of them (the targets) is highlighted in red, so as to obtain different load levels (i.e., 2, 3 or 4). Then, all objects turn black and start moving (b). Participants are instructed to track the target objects while they move. At the end of the tracking period (10 seconds), participants indicate which are the targets by clicking on them with the mouse (c). Crowding (i.e. spatial interference) was varied as the minimum distance at which distractors approached targets: >80 pixels in low crowding (e) and >15 pixels in high crowding (d).

The size of the display was 800×800 pixels (20.9 × 20.9 degrees of visual angle). Objects had a radius of 15 pixels (0.4 degrees). Trial duration was 10 seconds. The displays were redrawn at a rate of 30 frames per second and objects moved with a speed of 10 pixels per frame (0.27 degrees). We generated two sets of stimuli. In the “low crowding” trials the distance between targets and distractors that was always larger than 80 pixels (2.1 degrees; Figure 2.e). In the “high crowding” stimulus set, we allowed the distance between targets and distractors to be as short as 15 pixels (0.4 degrees; Figure 2.d). Close encounters in the high crowding trials were thus defined as situations where a distractor traveled within 15 and 60 pixels of a target (0.4 - 1.6 degrees) and kept a distance of less than 80 pixels for no more than 10 frames (333 ms). Thus, close encounters were brief, in order to ensure that some close encounters would not last over several seconds. Each load level in the “high crowding” set had exactly 13 close encounters, irrespective of load. In addition, it was ensured that each target had at least one close encounter with a distractor, but not more frequently than the 75% percentile of close encounter distributions in the generated trials. As close encounters could arguably still occur in the low crowding condition, we attempted to assert some control on the number of “long-range” close encounters. That is, we defined long-range close encounters as starting when a distractor approached a target closer than 100 pixels and ended when getting further away than 120 pixels. In selecting trials we ensured that the total number of long-range close encounters per trial would be less than the 75% percentile of such long-range close encounters in the generated set of trials prior to selection. Equalizing such long-range close encounters was not possible across load levels.

In summary, we had a mixed design with factors load (3 levels) and crowding (2 levels), making in total, six different conditions.

### 2.2 Eye-tracking study

#### 2.2.1 Participants

Forty-one adults (28 females) were recruited through social media to participate in the study (mean age: 24.2 years; range: 19-40 years). They were compensated with a gift card equivalent to 100 Norwegian Kroner (NOK) per hour and each session took approximately 1 hour. The study was conducted according to institutional guidelines, and was approved by the local ethics committee (IRB).

#### 2.2.2 Experimental procedure

The MOT task was written in code and delivered using custom software written in JavaScript. The participants read the instructions and performed a training session on a computer (10 trials). Afterwards, the full experiment was run in the lab. The total number of objects in the display was kept constant at 10. Each trial consisted of the presentation of the 10 objects in blue for 2 s. Then the target disks turned red for 2.5 s before turning back to blue. In the passive viewing condition, no ball turned red. The disks started to move and the tracking period lasted 10 s (tracking trials) or 7 s (passive viewing trials). At the end of the tracking period all objects’ movements stopped. Participants had to click on all the disks that were originally selected to be tracked. Participants clicked with the right button if they had high confidence on their response and with the left button if they had low confidence. Response accuracy and time to start responding were recorded for each trial.

#### 2.2.3 Pupillometry analysis

Blinks were removed and interpolated before hampel and lowess filtering were applied. For each blink detected by the SMI software, we linearly interpolated from 5 samples before onset until 5 samples after offset. Trials were baseline corrected using a baseline of 500 msecs before the beginning of the tracking period. The pupillometry data was analyzed in the time region 2633ms to 12000ms (determined by Monte Carlo analysis on load levels). To further explore whether the effect of crowding would become evident by the end of the trial, we analyzed the pupil change in two intervals, one early (2633 – 4633 msecs) and one late (10000 – 12000 msecs) within the trial. Saccades were detected with the SMI software and extracted during the tracking interval (2000-12000 ms).

### 2.3 fMRI study

#### 2.3.1 Participants

Forty adults (25 females) were recruited through social media to participate in the study (mean age: 25 years; range: 19-37 years). They were compensated with a gift card equivalent to 100 Norwegian Kroner (NOK) per hour and each session took approximately 2 hours.

Before starting testing, participants were asked to answer a brief questionnaire to assess whether they passed the inclusion criteria (no serious neurological or psychiatric illness, vision or language problems, etc.). Participants with an average proportion of accuracy of < 0.6 were removed since this level of performance approaches chance level. The study was conducted according to institutional guidelines and approved by the local ethics committee.

#### 2.3.2 Experimental procedure

The MOT task was included the same stimulus set as the eye-tracking study and was delivered using E-prime 2. The participants read the instructions and performed a training session before the fMRI session (10 trials). Different from the eye-tracking study, in the fMRI study we used partial report. At the end of the tracking period all objects’ movements stopped and one of the objects was highlighted with a yellow circle. The participant responded whether the probed object was one of the tracked objects. The response window was 2 s, after which the trial ended and was followed by a 4-s inter-trial interval (ITI) that consisted of a fixation cross. Response accuracy and reaction time were recorded for each trial. The probability that the probed object was one of the targets was 50% for all tracking conditions.

#### 2.3.3 fMRI acquisition and analysis

Participants were scanned in a 3T Philips MRI scanner at Rikshospitalet, Oslo. Each scanning session started with an anatomical scan (0.5 mm^3^). Four runs of functional images were acquired while participants performed the MOT task. One run consisted of 24 trials in two blocks of 12 trials. The load and crowding conditions were semi-randomized. A rest period of 20 s always followed after each block of trials. Whole-brain functional images were acquired using a spin-echo echoplanar (EPI) sequence sensitive to blood-oxygen-level-dependent (BOLD) magnetic susceptibility (TR = 2208 ms; Flip angle = 90deg; number of slices: 42; voxel size: 3 mm^3^). Each functional scan lasted about 8 minutes and in each scan, 234 volumes were collected. Finally, a neuromelanin-sensitive scan was acquired with the objective of further improving the localization of the locus coeruleus. The slices for this scan were set in order to cover the brainstem (T1-TSE, TR: 600ms, TE: 14ms, voxel dimension: 0.4×0.49×3mm, Flip angle = 90deg; number of slices: 10). This scan lasted 12 minutes.

The stimuli were projected onto a screen positioned at the head end of the litter. Participants viewed the screen through a mirror placed on the head coil. Participants’ response was given with the right hand through a joystick, with one button corresponding to “target” and another to “not a target” response.

The functional images of each participant were first visually inspected for anomalies and then submitted to a standard preprocessing pipeline using SPM 12 implemented on MATLAB (MathWorks, Natick, MA). The data of one participant could not be used because of errors saving the images. Images were first corrected for time delays and realigned using 6 parameters of movement. The data were normalized to a standard template and smoothed (8mm FWHM Gaussian kernel). For the analysis of the activity in the locus coeruleus, a smaller Gaussian kernel was used (3mm FWHM). Additional motion correction was performed by scrubbing the volumes with excessive movement using FSL functions.

Preprocessed images were submitted to a first level analysis. Event-related activation was estimated with a general linear model (GLM). Stimulus presentation and tracking intervals of the different load and crowding levels were modeled as separate events with a canonical HRF. Images were high-pass filtered at 128 sec. Contrasts were generated for each tracking condition both collapsing across load levels and for each load level. Parameter estimates from each participant’s GLM were submitted to a second-level test. A full-factorial analysis was performed treating participants as a random factor, and load and crowding as the fixed factors. The maps of load 4 > load 2 and high > low crowding were masked by the respective (load or crowding) main effects. To obtain distinct areas in load and crowding, each of the former maps was masked by the other using thresholds of p = 0.05 and p = 0.001 for the exclusive mask and the contrast, respectively.

Given our interest in the brainstem, we separately assessed the activity in the VTA/SN and LC. For the VTA/SN, we performed a small volume correction in the contrasts of interest on a combined VTA/SN mask derived from probabilistic atlases of the structures (Murty et al., 2014). The VTA and SN masks partially overlap and therefore we combined them; the resulting mask was cropped to be spatially restricted to the brainstem. For the LC, given its small size and the variability of its location in the brainstem across individuals, we defined individual masks in the high-resolution T1-TSE images and used them for a region of interest analysis. Specifically, the LC of each individual was delineated on the axial slices of the neuromelanin scans. The position of the nuclei was determined in the pons as the voxels of hyperintensity on either side of the fourth ventricle, following the procedure employed in previous studies (Krebs et al., 2018). The T1-TSE scan of each individual was coregistered to the corresponding structural image, the structural was coregistered to the mean functional and then the deformation field calculated during the normalization step was applied to the coregistered T1-TSE image. Region of interest (ROI) analyses were performed on the individual masks of the locus coeruleus. The data were extracted from the contrast images with rfxplot toolbox (Gläscher, 2009). The parameter estimates in the voxel of maximum effect size in each condition were extracted for the different load levels within the masks. Then we performed RM-ANOVA applying load (levels: 2, 3 and 4 objects) and crowding (low, high) as within-subject factors. For the comparison of groups, we included group (high-performers and low-performers) as between-subjects factor. For the comparison of task-related activity in the different groups in the VTA, a similar ROI analysis was applied on the VTA/SN combined mask.

### 2.4 Data analysis: behavior

Errors, reaction times, confidence estimates, pupil change, number of saccades and brainstem activity for each condition were compared with repeated measures (RM) ANOVA, correcting for non-sphericity. Within-subject factors were load (3 levels) and crowding (2 levels). Only correct trials were included in the RT analysis. For the analysis of the groups of performers, we divided the participants into high and low performers on the basis of a median split of total tracking accuracy. We performed RM-ANOVA as before adding group as a between subject factor. For this analysis, only the lowest and highest load levels were considered as to look at the largest effects. All effect sizes are reported as generalized eta squared (ηG^2^). The data was analyzed with IBM SPSS.

## 3. Results

### 3.1 Eye-tracking study

#### 3.1.1 Behavior

Increasing load caused a moderate decrease in accuracy (F(1.92,76.67) = 20.92, p < 0.001, η_G_^2^ = 0.07, Figure 3). Furthermore, high crowding also caused a decrease in accuracy (F(1,40) = 510.56, p < 0.001, η_G_^2^ = 0.54), and interacted weakly with load (F(1.68,67.22) = 5.99, p = 0.006, η_G_^2^ = 0.01). Confidence estimates were calculated for each trial as the percentage of responses that were given with the left mouse button (high confidence). An RM-ANOVA on confidence estimates showed significant main effects of load (F(1.39, 55.52) = 25.27, p < .0001, η_G_^2^ = 07) and crowding (F(1, 40) = 50.64, p < .0001, η_G_^2^ = .18), with lower confidence at higher difficulty in both effects. The interaction was not significant (F(1.69, 67.51) = 3.29, p = 0.051, η_G_^2^ = .003).

**Figure 3.**
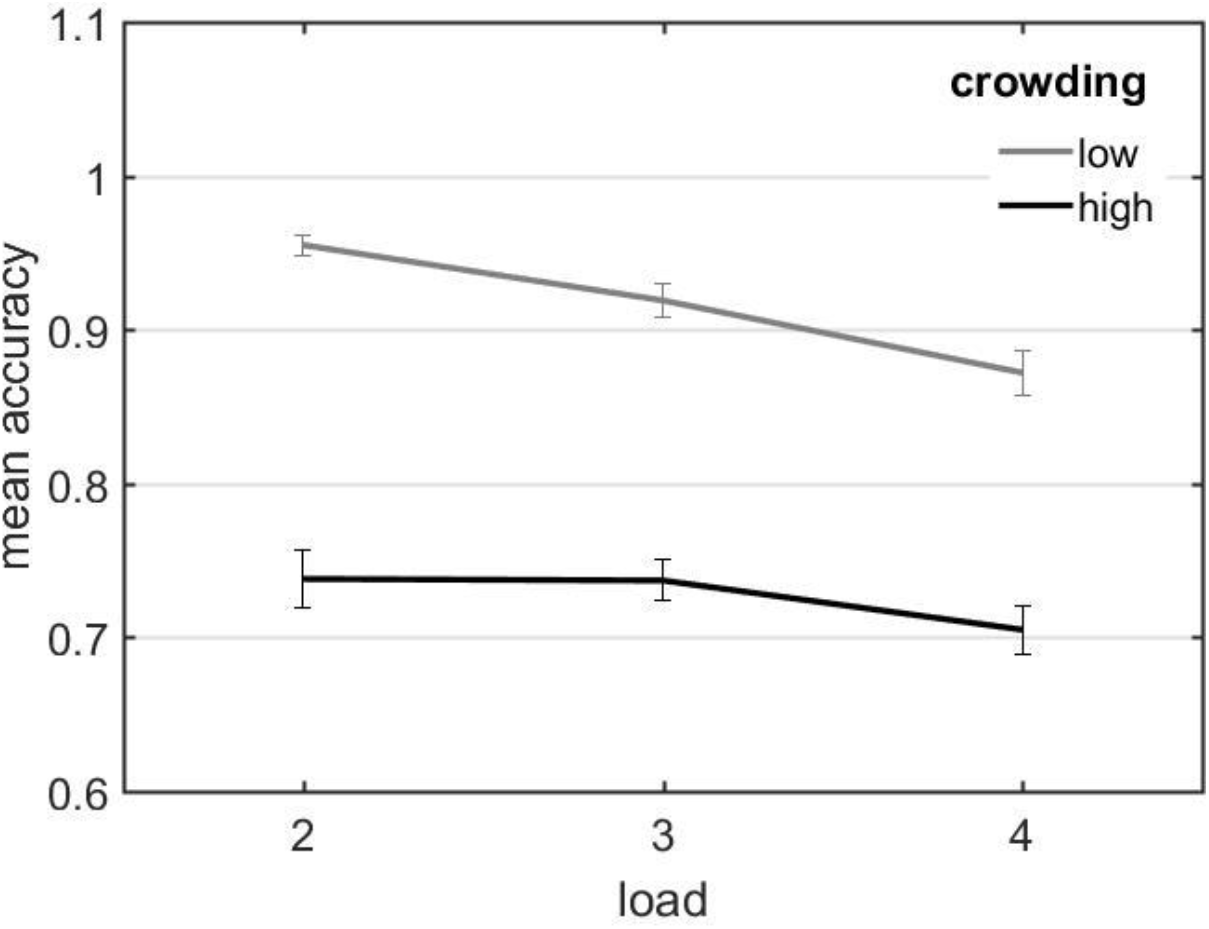
Behavioral results of pupillometry study. Mean accuracy as a function of load for each of the crowding levels.

#### 3.1.2 Pupil response

We analyzed the effects of load and crowding on pupil dilation during tracking. We found a significant increase in pupil size with load (main effect of load: F(1.77,70.86) = 57.6, p < 0.001, η_G_^2^ = 0.04; Figure 4). Crowding (F(1,40) = 0.08, p = 0.78), or the interaction between load and crowding (F(1.96, 78.25) = 1.26, p = 0.29), did not show significant effects.

**Figure 4.**
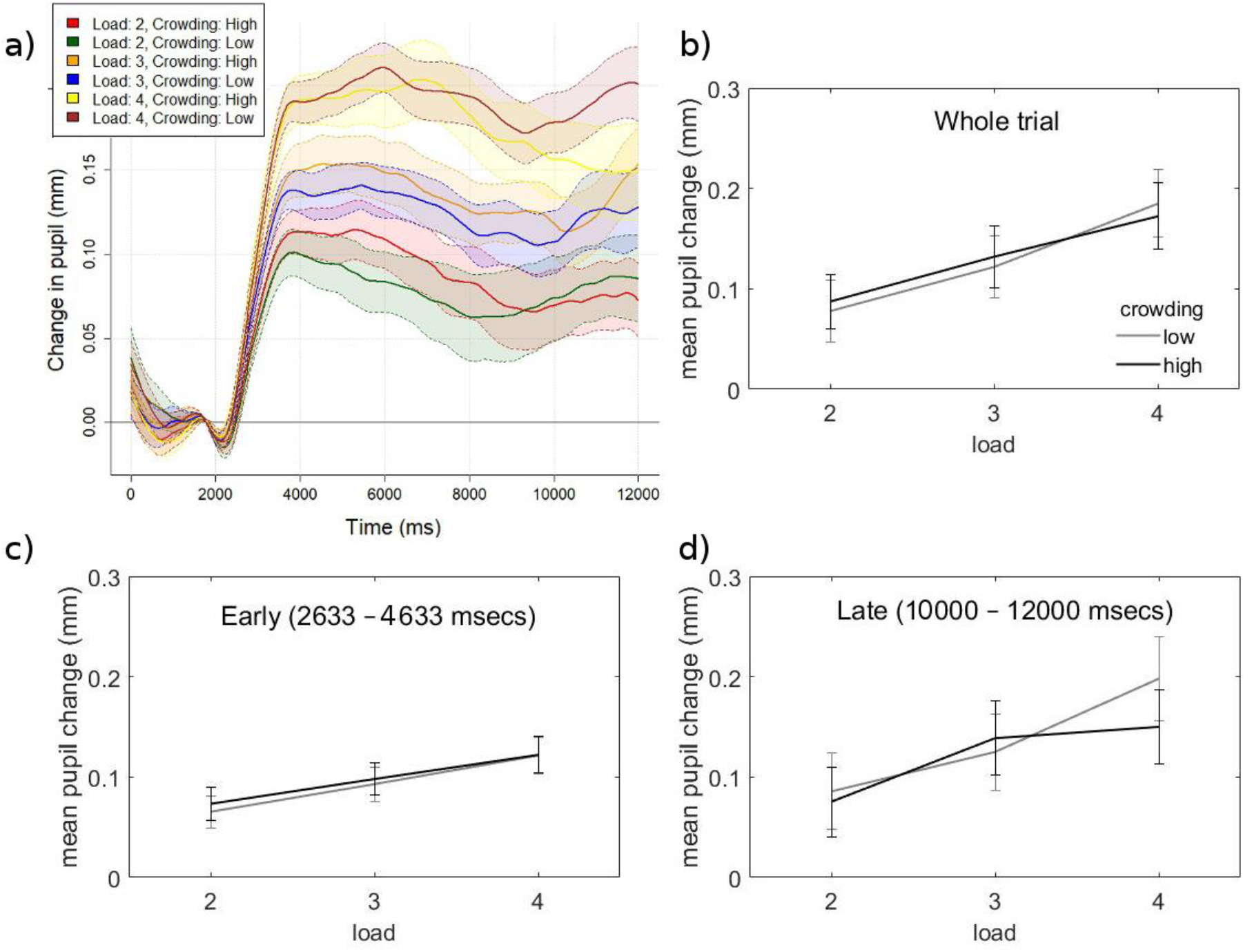
Evoked pupil size during tracking for the different conditions. a) Baseline corrected change in pupil during tracking at the indicated conditions. Shaded are indicates standard error. b-d) Average pupil changes across the whole trial (a), an early interval (b) or a late interval (c) within a trial. Error bars indicate the standard error of the mean.

When analyzing the pupil change early or late within the trial, the effect of load was significant in both intervals (Early: F(1.93, 77.25) = 48.73, p < 0.001, η_G_^2^ = 0.04; Late: F(1.83, 73.37) = 34.27, p < 0.001, η_G_^2^ = 0.02; Figure 4). The effect of crowding or the interaction was non-significant in any interval (p > 0.28).

#### 3.1.3 Saccade frequencies

For this analysis, we removed two participants for having an unusual high number of saccades (more than an average of 25 saccades per trial). Number of saccades showed a significant effect of load (F(1.59, 60.56) = 7.01, p = 0.004, η_G_^2^ 0.004), and crowding (F(1, 38) = 34.88, p < 0.001, η_G_^2^ = .02). The interaction was also significant (F(2.00, 75.95) = 3.18, p = 0.047, η_G_^2^ = 0.0006). On average, participants made 11.1 saccades in high crowding trials and 9.2 saccades in low crowding trials.

#### 3.1.4 Individual differences

We inspected the differences in behavior and ocular measures between high and low performers due to load and crowding (Figure 5). Accuracy presented a significant interaction between group and load (F(1.95, 75.94) = 3.85, p = 0.030, η_G_^2^ = 0.03) and group and difficulty (F(1,39) = 27.65, p < 0.001, η_G_^2^ = 0.08) and three-way interaction (F(1.85, 72.2) = 10.69, p < 0.001, η_G_^2^ = 0.04). In low crowding, both groups had significantly lower accuracy with load (high performers: t(20) = 3.04, p = 0.006; low performers: t(19) = 9.56, p < 0.001). In high crowding, high performers had lower accuracy with load (t(20) = 2.16, p = 0.043) while low performers showed no differences with load (t(19) = 1.15, p = 0.262).

**Figure 5.**
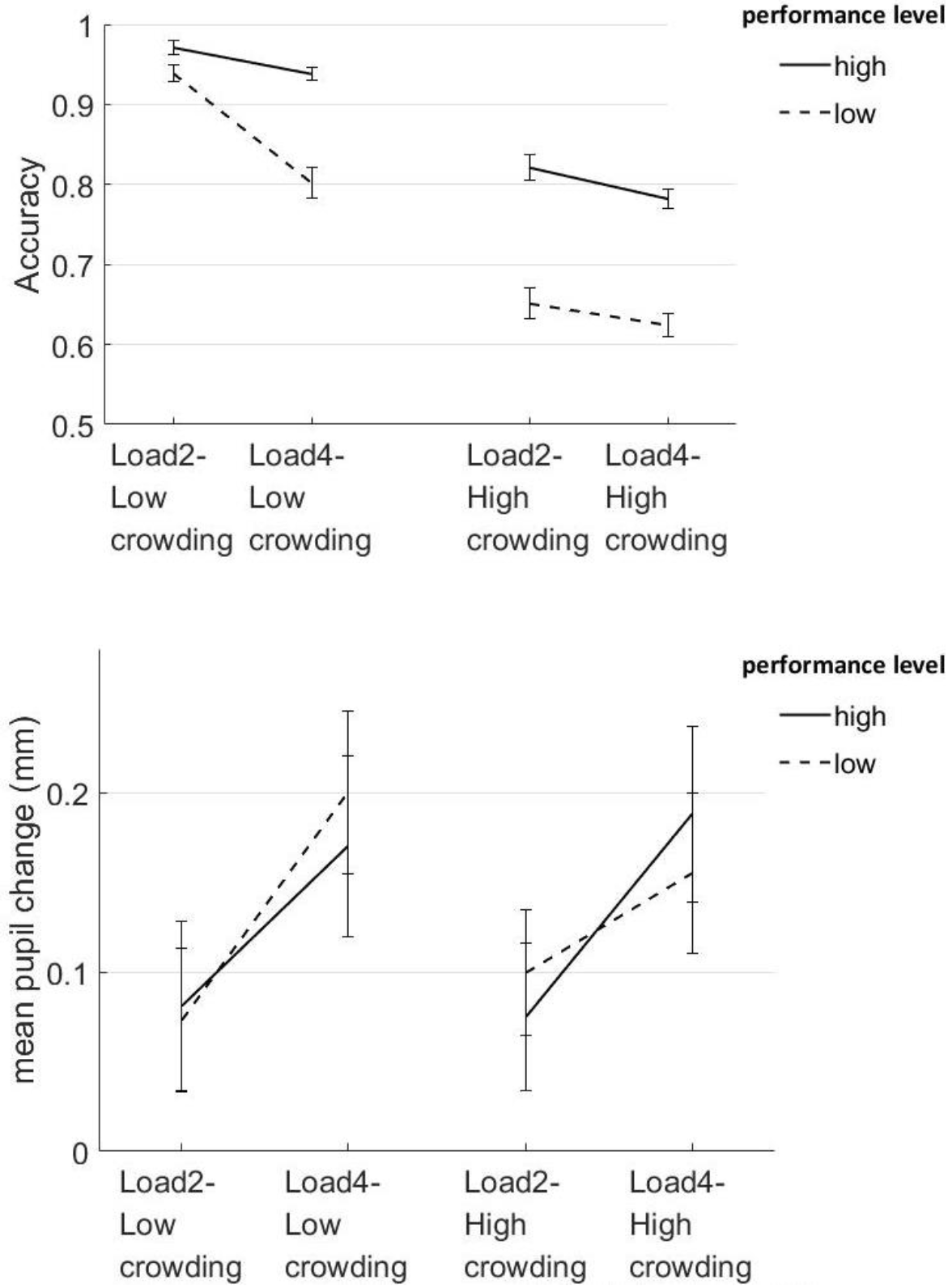
Behavioral and pupillometry results for performers groups. Mean accuracy (top panel) and mean pupil response (bottom panel) as a function of load for each of the crowding levels and groups.

The analysis of group differences on evoked pupil revealed a significant effect of load (F(1.77, 69.18) = 56.24, p < 0.001, η_G_^2^ = 0.04), and an interaction between performer group, load and difficulty (F(1.99, 77.45) = 5.17, p = 0.008, η_G_^2^ = .003). In load 4, low performers had significantly less pupil dilation in high crowding compared to low crowding (t(19) = 3.31, p = 0.003), but not high performers (p = 0.31).

Next, we analyzed saccade frequencies between high and low performers. High performers made an average of 8.7 saccades per trial, while low performers made an average of 11.9 saccades per trial, this difference was however not statistically significant (t(25.9) = 1.5, p = 0.14). Both of the removed participants were classified as low performers.

### 3.2 fMRI study

#### 3.2.1 Behavior

Increasing load caused a moderate decrease in accuracy (F(2,76) = 11.98, p < 0.001, η_G_^2^ = 0.08, Figure 6). Furthermore, high crowding caused a large decrease in accuracy (F(1,38) = 114.61, p < 0.001, η_G_^2^ = 0.25), and did not interact with load (F(2,76)= 0.40, p = 0.63). RT increased with higher load (F(2,76) = 4.42, p = 0.015, η_G_^2^ = 0.01) and higher crowding (F(1,38) = 25.85, p < 0.001, η_G_^2^ = 0.02). There was no interaction between load and crowding in RTs (F(2,76)= 0.82, p = 0.42).

**Figure 6.**
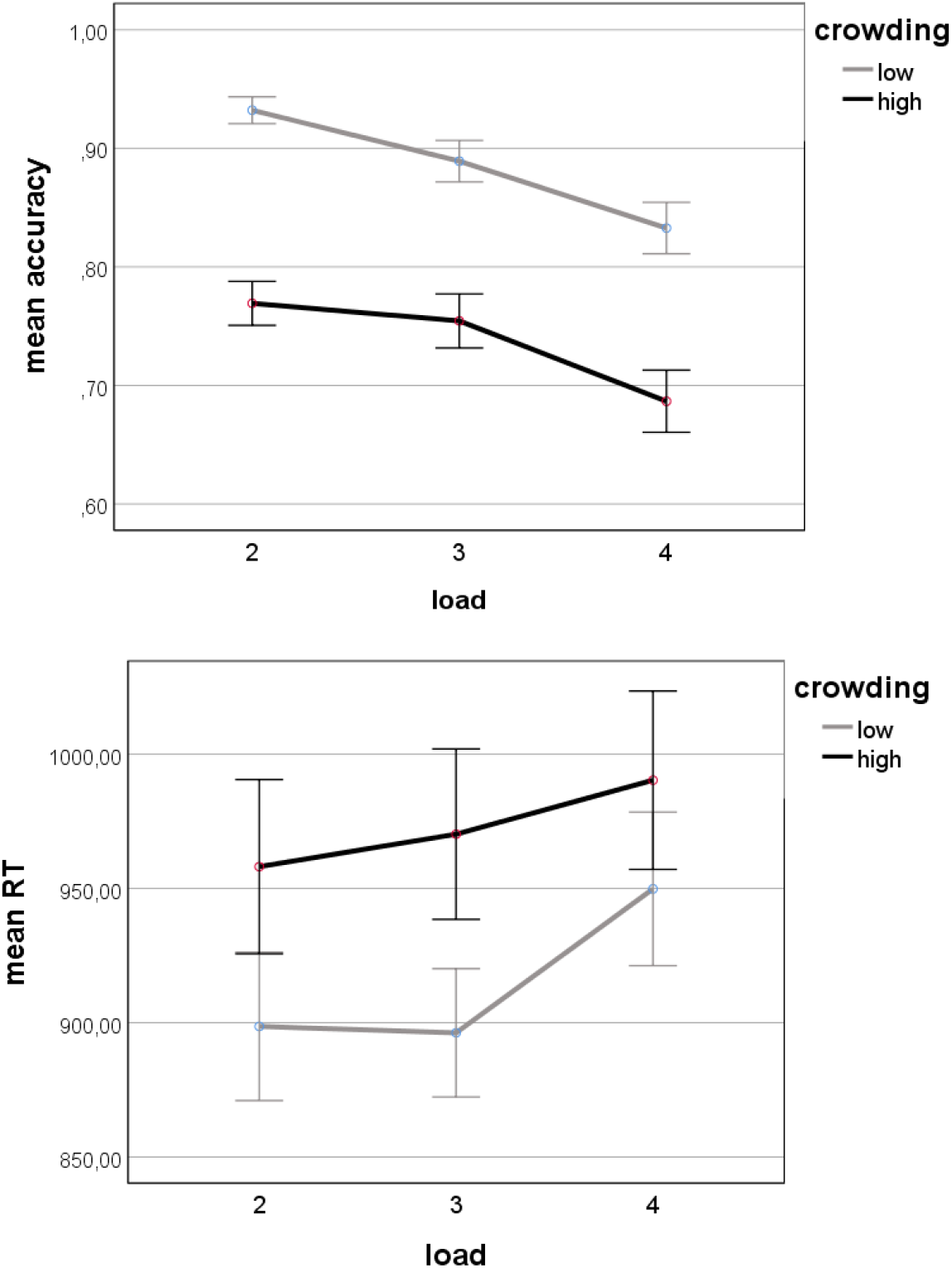
Behavioral results of fMRI study. Mean accuracy (top panel) and RT (bottom panel) as a function of load and for each crowding level. Error bars indicate the standard error of the mean.

#### 3.2.2 Whole brain fMRI analysis

A set of areas was associated to tracking as compared to passive viewing, comprising the bilateral inferior/middle occipital cortex, the bilateral inferior/superior parietal cortex and bilateral middle frontal cortex, precentral area and supplementary motor area. (Figure 7, Tables 1 and 2). This pattern replicates previous findings on MOT and fMRI (Culham et al., 1998).

**Figure 7.**
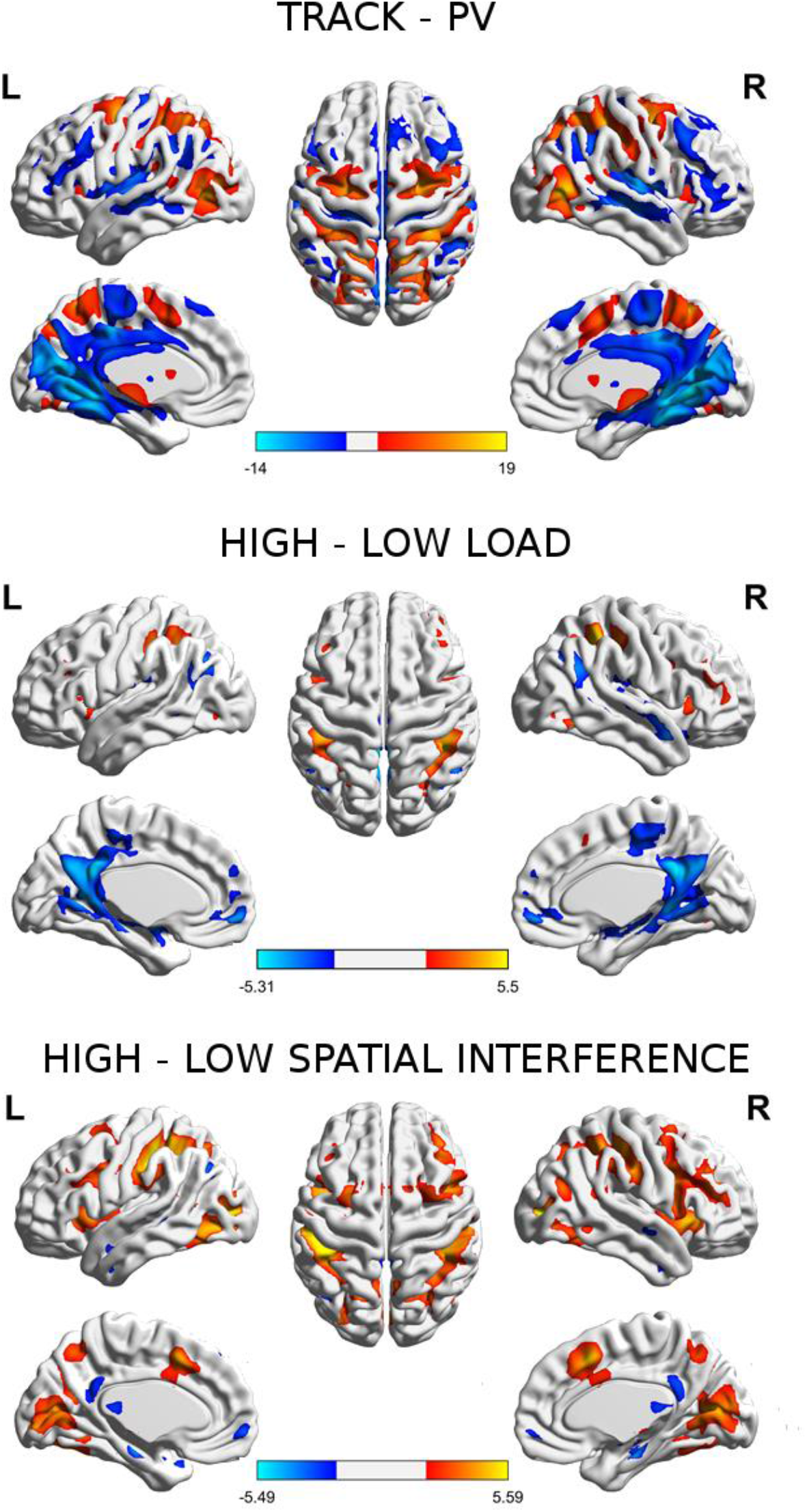
Brain regions’ activations and deactivations obtained from the indicated contrasts. L: left hemisphere; R: right hemisphere. The colorbar indicates the obtained t value.

**Table 1.** Summary of brain areas presenting hemodynamic changes due to the different conditions: Tracking > passive viewing (Tracking), Load 4 > Load 2 (Load), and High > Low crowding (Crowding). Positive activations are marked with a plus sign (+), and negative with a minus sign (-). If activations were lateralized, they are indicated as L = left, R = right. For precise coordinates, please refer to Tables 2 and 3.

|  | Tracking | Load | Crowding |
| --- | --- | --- | --- |
| **Frontal** |  |  |  |
| Supplementary motor area | + |  | + |
| Precentral gyrus | + |  | + |
| Middle frontal gyrus (including Frontal eye fields) | + |  | + ( L ) |
| Middle frontal gyrus | + | + ( R ) | + ( R ) |
| Inferior frontal gyrus |  | + ( L ) |  |
| Medial frontal gyrus |  |  | + |
| Frontal medial orbital |  | - | - |
| Anterior Insula |  |  | + |
| Posterior insula | - |  |  |
| **Parietal** |  |  |  |
| Superior parietal lobule | + |  |  |
| Inferior parietal lobule | + | + | + |
| Middle/posterior cingulate cortex | - | - |  |
| **Temporal/occipital** |  |  |  |
| Fusiform |  | + | + |
| Middle/superior temporal gyrus | - | - |  |
| Precuneus | + |  |  |
| Inferior occipital gyrus | + |  |  |
| Middle occipital gyrus | + |  | + |
| Lingual gyrus |  |  | + |
| Calcarine | - | - |  |
| **Subcortical** |  |  |  |
| Thalamus (pulvinar) | + |  |  |
| Midbrain | + |  |  |
| Hippocampus |  | - | - |

Load evoked parametrically higher activation in bilateral inferior parietal area and left inferior frontal (Figure 7, Tables 1 and 3). High crowding evoked higher activity in middle occipital areas, as well as bilateral inferior parietal and bilateral middle frontal gyrus/precentral area including frontal eye fields, and insula (Figure 7, Tables 1 and 3). The difference between these contrasts revealed that crowding presented additional activity in bilateral inferior parietal and medial frontal regions, insula and primary visual areas (lingual/calcarine) as compared to load (Figure 8, Table 3).

**Figure 8.**
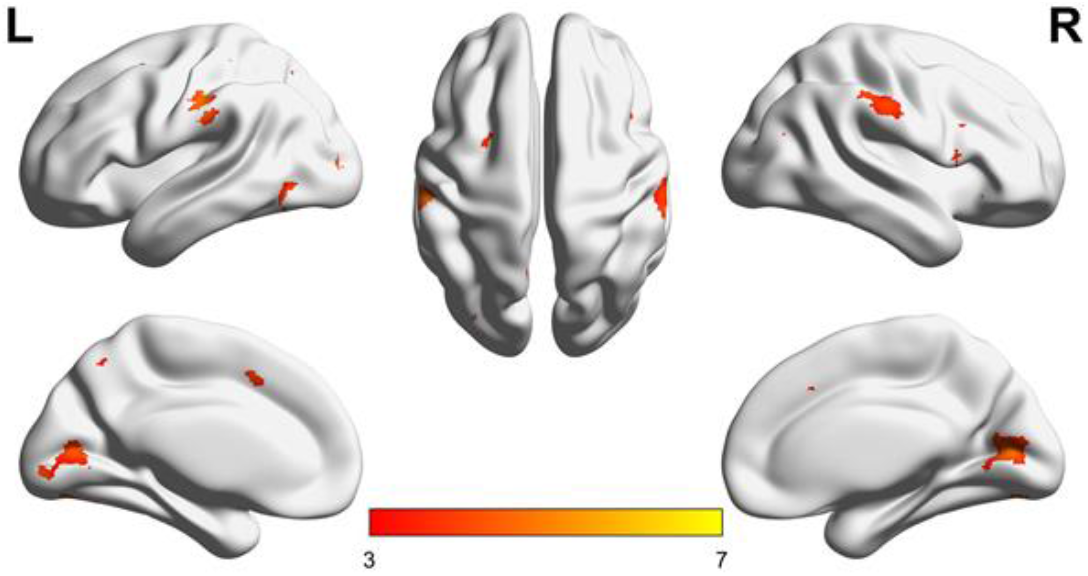
Effect of high – low crowding (masked Load 4 – load 2)

**Table 2.**

| Track-PV |  |  | cluster |  | Coordinates |  |  |
| --- | --- | --- | --- | --- | --- | --- | --- |
| Hemisphere | Area | BA | p | size | x, y, z | peak p | T |
| RH | Middle Occipital Gyrus | BA 37 | <0.01 | 16399 | 46 -68 4 | <0.001 | 19.38 |
| LH | Inferior Occipital Gyrus | BA 19 |  |  | -44 -76 4 | <0.001 | 18.17 |
| RH | Precuneus | BA 7/31 |  |  | 36 -40 54 | <0.001 | 15.07 |
| LH | Superior Parietal Lobule | BA 7 |  |  | -24 -60 58 | <0.001 | 13.64 |
| RH | Inferior Parietal Lobule | BA 40 |  |  | 42 -30 42 | <0.001 | 11.00 |
| RH | Middle Frontal Gyrus | BA 6 | <0.01 | 6445 | 24 -6 52 | <0.001 | 17.67 |
| LH | Middle Frontal Gyrus | BA 6 |  |  | -24 -6 60 | <0.001 | 13.93 |
| RH | Precentral Gyrus | BA 6 |  |  | 56 6 32 | <0.001 | 9.63 |
| RH | SMA | BA 6 |  |  | 8 6 52 | <0.001 | 8.42 |
| LH | Precentral Gyrus | BA 6 |  |  | -40 -8 54 | <0.001 | 7.75 |
| RH | Midbrain |  | <0.01 | 1253 | 4 -18 -6 | <0.001 | 7.56 |
| RH | Thalamus | Pulvinar | 0.161 | 131 | 14 -24 14 | <0.001 | 5.95 |

**Table 3.**
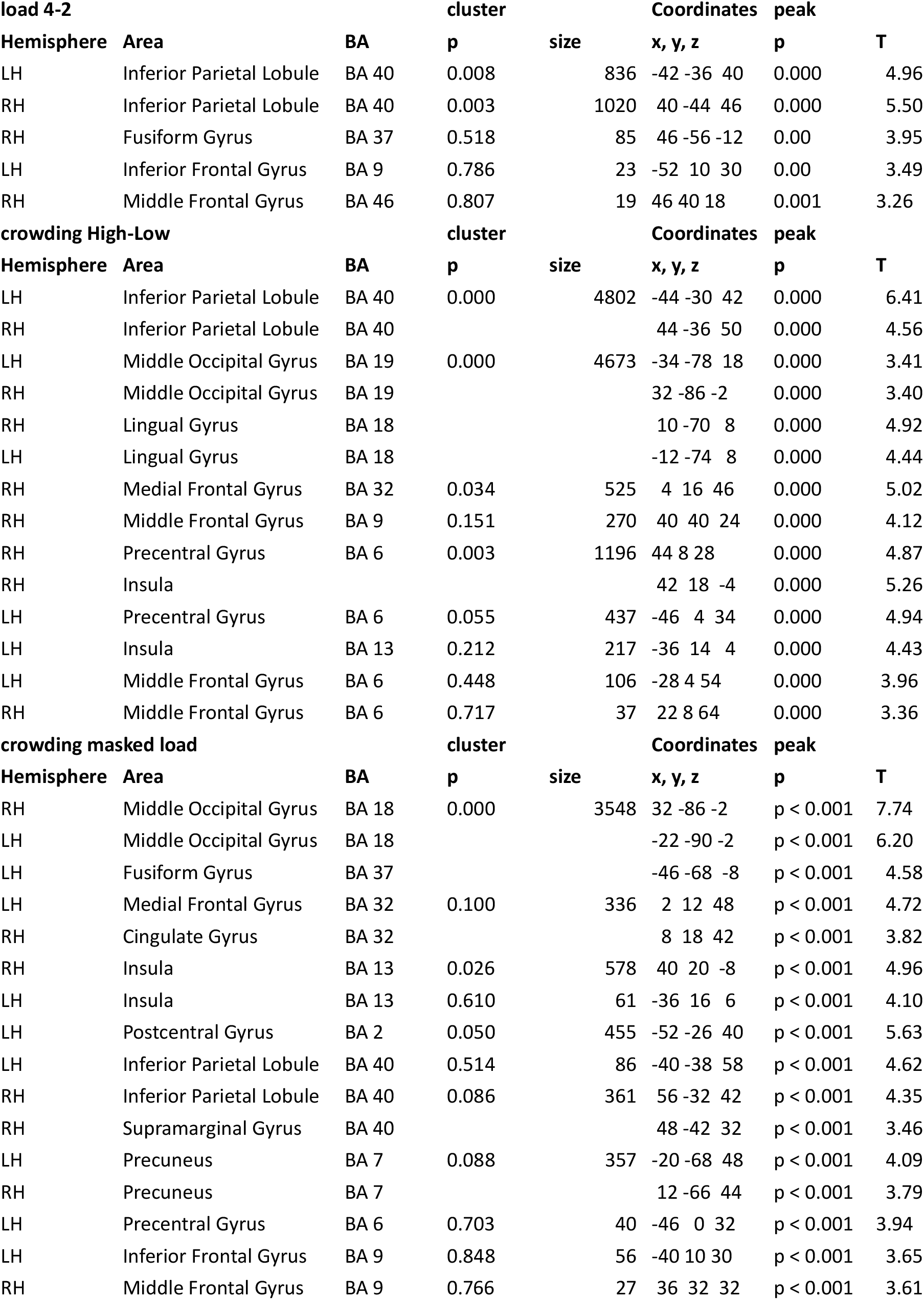
Brain coordinates of activation at the indicated contrasts

#### 3.2.3 Brainstem nuclei

We inspected the effect of increasing effort on the activity within individually defined LC masks. We found a significant effect of load (F(2,74) = 3.21, p = 0.046, η_G_^2^ = 0.02, Figure 9). We found no effect of crowding (F(1,37) = 0.59, p = 0.44) or interaction (F(2,74) = 3.21, p = 0.066).

**Figure 9.**
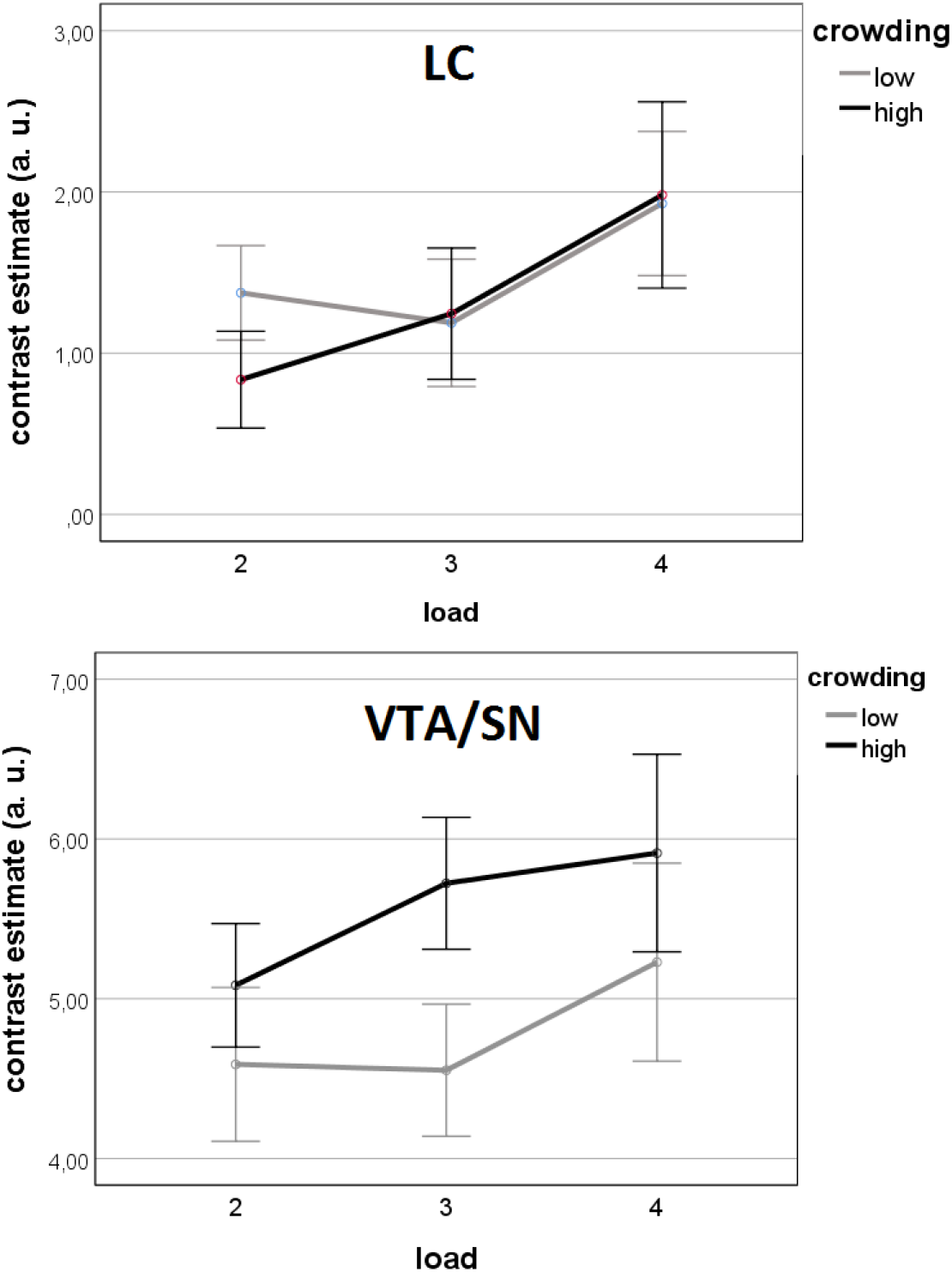
Brainstem activity evoked by the different conditions. Top: LC; bottom: VTA/SN

Moreover, load had no significant effect on activity in the VTA/SN nuclei (F(2,74) = 1.71, p = 0.194, Figure 9) but high crowding evoked a larger activity than low (F(1,37) = 4.96, p = 0.032, η_G_^2^ = 0.02). We found no interaction (F(2,74) = 0.375, p = 0.656).

#### 3.2.4 Individual differences

High and low performers showed different levels of accuracy for the different conditions. In addition to the main effects of load and crowding (F(1,36) = 28.51, p < 0.001 η_G_^2^ = 0.11 and F(1,36) = 88.14, p < 0.001, η_G_^2^ = 0.27, resp., Figure 10), we found a significant interaction of performance level by load (F(1,36) = 9.20, p = 0.004, η_G_^2^ = 0.04). Low performers had lower accuracy in load 4 as compared to load 2 (T(19) = 5.50, p < 0.001), while the effect of load in high performers did not reach significance (T(17) = 1.83, p = 0.084). No group effects were observed for RT.

**Figure 10.**
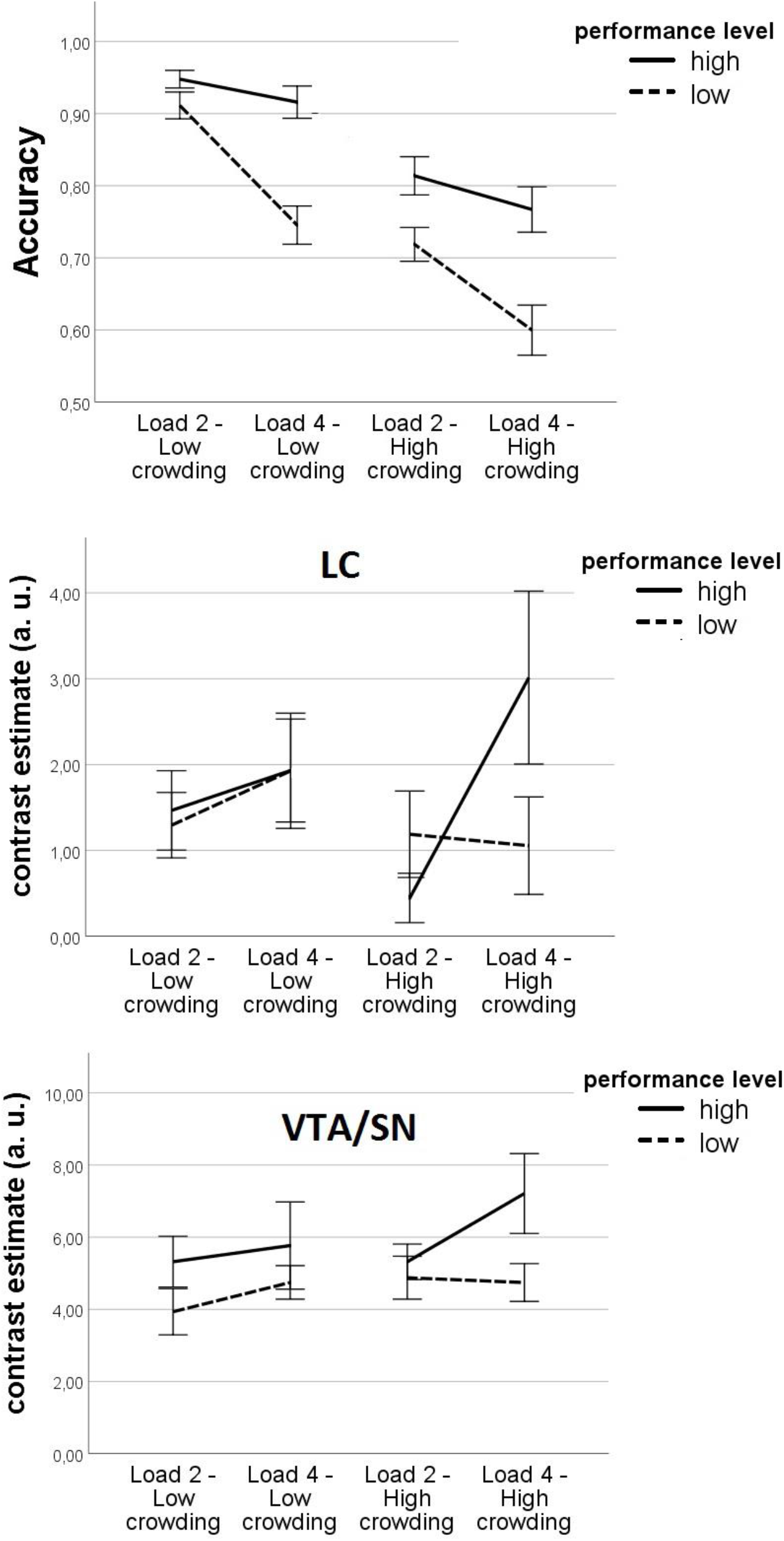
Behavioral and brainstem effects on high and low performers. Top panel: Mean accuracy; middle panel: LC; bottom panel: VTA/SN, in the high and low crowding conditions for the lowest and highest load level split by high performers versus low performers.

In the LC, we found a significant three-way interaction of load by performance level by crowding (F (1,36) = 4.44, p = 0.042, η_G_^2^ = 0.02, Figure 10). When looking separately at the different crowding levels, we observed a significant load by group interaction (F(1,36) = 5.55, p = 0.024, η_G_^2^ = 0.06) in the high but not in the low crowding condition. With high crowding, high performers had higher activity in load 4 as compared to load 2 (T(17) = 2.51, p = 0.022), but not low performers (p > 0.2 for all contrasts).

In the VTA/SN, we found no main effect or interaction effects (Figure 10). High performers presented larger activity than low performers, although the results were not significant (F(1,36) = 3.17, p = 0.083).

## 4. Discussion

Based on the convergent findings obtained from our two independent studies, we conclude that load and crowding are handled by distinct processes that rely, at least partially, on different neural mechanisms. This conclusion is based on the observation that, even when the level of spatial interference (low/high crowding) was kept equal, accuracy decreased and RT increased with load level (number of targets), hinting that crowding and load pose challenges to brain processing differentially. In line with this, increasing demands due to load were signaled by increasing arousal (as indicated by pupil size) and LC activity across the tracking period. Increased load elicited proportionally greater activation in the fronto-parietal attention network and more frequent saccades. In addition, crowding events engaged the VTA/SN, and further implicated the attentional network and number of saccades. Furthermore, high performers appeared able to recruit their cognitive resources during stimulus-driven conflict, as signaled by higher pupil and LC activity during high crowding. Taken together, the above results suggest that the brain can cope with spatial interference and effort in a highly specific and adaptive manner.

### 4.1 Behavioral effects of load and spatial interference

We note that different factors are known to affect performance in MOT: number of objects or cognitive workload (both targets and non-targets), crowding, speed and hemifield (S. L. Franconeri et al., 2013; Meyerhoff et al., 2017; Scimeca & Franconeri, 2015). All these factors seem to share a common property: they affect the degree and frequency of proximity between objects. Hence, a still open question regarding the MOT paradigm was whether the deterioration of performance could be simply reduced to spatial interference, that is, to a perceptual limitation in attentional resolution. For example, increasing number of targets could imply that more close encounters with other objects will likely occur and, therefore, load should correlate with number of close encounters. Some studies have assessed the effect of inter-object spacing on accuracy, either at different or at a fixed level of load (Alvarez & Franconeri, 2007; Bae & Flombaum, 2012; Feria, 2013; S. L. Franconeri et al., 2008; Iordanescu et al., 2009; Shim et al., 2008). Others have assessed the relation between load and speed (S. Franconeri et al., 2010; Srivastava & Vul, 2016) or between speed and object spacing (Alvarez & Franconeri, 2007; Vul et al., 2009). In the current study, we systematically varied load at two levels of crowding. This allowed us to assess the relation between the factors.

A decreasing linear trend in accuracy with load at both levels of crowding suggests that, in addition to the frequency of close encounters, the number of objects pose an additional demand on the participants, and therefore those factors rely on partly non-overlapping mechanisms (Figure 1.b). Given that the spatial interference was equalized within crowding levels and across load demands, our results indicate that load may pose an additional demand on attentional functions. This agrees with previous work suggesting that recruiting higher order mechanisms allow for dynamic allocation of attention to crowded situations (Intriligator & Cavanagh, 2001; Iordanescu et al., 2009; Z. Pylyshyn, 2004; Srivastava & Vul, 2016). Such higher order mechanisms likely involve visual selection and sustained visual attention, which may be specifically engaged during target assignment or during the tracking period, respectively (Meyerhoff et al., 2017). While the above terms may seem somehow loosely defined mechanistically, it has been shown that cognitive processes, such as update in working memory, are engaged at the assignment of objects (Allen et al., 2006; Ma & Flombaum, 2013) and during tracking (Eayrs & Lavie, 2018; Oksama & Hyönä, 2004), consistent with computational simulations (Srivastava & Vul, 2016; Vul et al., 2009). Similarly, Vul et al. (2009) argued that “while many results in MOT arise as consequences of the information available for the computational task, the speed-number tradeoff seems to be the result of a flexibly-allocated resource such as memory or attention”. Possibly, cognitive processes may help to circumvent the limitations of the visual attentional processes and vice versa (Oksama & Hyönä, 2004). In line with this, a series of EEG studies have shown that different ERP signatures were related to the demands exerted by target number and the demands of the tracking itself (Drew et al., 2013; Drew et al., 2009; Drew & Vogel, 2008). For example, a parieto-central negativity was related to increasing target number during the assignment period, a contralateral delay activity was related to load during the tracking period, and early negative and positive components were related to following targets rather than distractors during the tracking period. These results argue for a highly coordinated and dynamic process developing during the tracking to keep target information in working memory, resolve confusion and remain engaged.

### 4.2 Pupil size increases with load but not with crowding

We replicated the finding that pupil size is sensitive to load (Alnæs et al., 2014; Wright et al., 2013). One possibility is that pupil size reflects preparatory effort mechanisms recruited in the beginning of the trial and sustained during the whole trial. This would agree with the proposal that pupil dilation reflects a readiness to expend resources (Bruya & Tang, 2018; van der Wel & van Steenbergen, 2018). Importantly, we found that pupil size was not sensitive to our manipulation of spatial interference. Thus, one could argue that the effects of load and crowding rely on different mechanisms, for example different neural mechanisms (as discussed below). We also note that effort due to the load manipulation is sustained over the tracking period, whereas the close encounters are transient events. It was expected that the effect of such transient events would accumulate to such an extent that they would influence the average pupil size over the entire tracking period, but it is also possible that these effects would wash out in the average.

The number of saccades increased with load and crowding, although in a greater extent with crowding. This result may correspond to the increase in activity that we observed in the frontal eye fields (FEF) in high versus low crowding. FEF are involved in the control of eye movement (Vernet et al., 2014) and may have being implicated in directing the eyes towards highly conflicting events (i.e. close encounters), what is consistent with previous results on MOT and eye-tracking (Zelinsky & Todor, 2010).

### 4.3 Cortical attentional networks related to both load and spatial interference

We found that a fronto-parietal network including inferior parietal and dorsal frontal areas, as well as occipital areas, were recruited during tracking. Load was further related to increased activity in the bilateral inferior parietal and inferior frontal areas. These activations closely match the ones reported by previous studies on MOT (Alnæs et al., 2014; Culham et al., 1998; Culham et al., 2001; Jahn et al., 2012; Jovicich et al., 2001; Shim et al., 2009; Tomasi et al., 2004). The fronto-parietal network is generally activated during demanding visuo-spatial tasks of different nature (Corbetta & Shulman, 2002; Duncan, 2010; Hugdahl et al., 2015), suggesting that its components cooperate to represent voluntary and stimulus-driven attentional priority (Ptak, 2012; Scolari et al., 2015; Serences & Yantis, 2006). As part of this network, the inferior parietal cortex is suggested to index the location of selected or prioritized objects (Serences & Yantis, 2006) and its activity depends on capacity limitations in visual short-term memory (Mitchell & Cusack, 2007). The inferior parietal cortices interact with superior parietal and frontal areas (Howe et al., 2009) and constitute the so-called dorsal attention network (Corbetta & Shulman, 2002; Fox et al., 2006; Vossel et al., 2014), whose involvement is consistent with top-down processes engaged during the task to guide the tracking of the target objects.

Higher crowding presented strong activity in bilateral inferior parietal, dorsal-frontal, including FEF, and occipital regions. In addition, insula and medial frontal areas were engaged. The insula and medial frontal areas are considered parts of the ventral attention network (Corbetta et al., 2008; Vossel et al., 2014), whose function is thought to be detecting salient events and assist attention orienting. During tracking, events such as close encounters may engage the ventral attention network to flexibly allocate attention where it is most needed. Therefore, an increased activity in high vs. low crowding may be related to coping with the increasing on-line demands imposed by that manipulation.

The frontal eye fields were found to be activated during high versus low crowding. In a seminal study on MOT with fMRI, Culham et al. (1998) found that the activity in FEF was related to tracking but was not modulated by load. Here we corroborate this finding, and in addition, we propose that the FEF activity during tracking is related to the crowding events, which were not controlled for in Culham et al. study. Therefore, its involvement in the task seems to be active in the resolution of moment-to-moment confusion, rather than barely “suppressing eye movements” (Culham et al., 1998).

Previous studies showed specificity in the factors accounting for the activation of some areas during multiple-object tracking (Atmaca et al., 2013; Howe et al., 2009; Merkel et al., 2015). Here we show that large activations in attentional and occipital areas were observed due to spatial interference as compared to load. Strong activation in the medial superior parietal lobe (including precuneus) has been suggested to reflect voluntary shifts of attention between perceptual entities (Serences, Liu, et al., 2005; Serences, Shomstein, et al., 2005; Serences & Yantis, 2006; Yantis et al., 2002). Other areas, such as medial frontal cortex and insula, were recruited additionally during high vs. low crowding and as compared to the load effect. A note of caution should be made given that it is not possible to know whether load 4, compared to load 2, imposed as high demands as high vs. low crowding, as indicated by the different effect sizes of those two factors on accuracy. Indeed, previous studies have shown larger brain effects of load when including load 5 (e.g. (Alnæs et al., 2014)). Therefore, we cannot exclude that the network of cortical areas underlying load and crowding were the same and that differences in the demand levels of our paradigm caused that the effects of crowding in brain activity to be stronger than the effect of load.

### 4.4 Brainstem neuromodulatory systems

Neuromodulators play an active role on attention by adjusting responsivity of neurons to optimize processing of inputs, a concept known as “gain” (Aston-Jones & Cohen, 2005; Servan-Schreiber et al., 1990). In the context of the MOT, we found that two nuclei, locus coeruleus (noradrenergic) and VTA/SN (dopaminergic), presented different profiles for the different task manipulations. LC increased activity with load, while VTA/SN showed increased activity with spatial interference. The involvement of LC agrees with the findings from a previous study which employed an atlas-based mask of the LC to extract the task-related activity (Alnæs et al., 2014). The LC/NE system has been studied in the context of mental effort and its function has been posited to be to increase the gain of the neural circuits relating co-occurring events. This result may be explained by the recruitment of LC by a top-down frontal system engaged in the preparation of load-related effort. We further found that LC activity was not significantly related to spatial interference. In contrast, the VTA/SN presented higher activity in high with respect to low spatial interference, but no effect with cognitive workload. Such a double dissociation between brainstem systems is (to the best of our knowledge) novel within human imaging studies in the context of an attentional task with varying levels of cognitive effort. Interestingly, the VTA/SN and LC share reciprocal connections (Weinshenker & Schroeder, 2007), and therefore they are expected to be tightly related. Within theories of cognitive effort, LC activity may be recruited after the computation of cost at the initiation of the trial, when the targets’ number or load is known. VTA/SN activity may be, on the other hand, related to working memory processes that enable propagation of state estimates of the position of the targets through time. In other words, one mechanism (VTA/SN) may depend on events occurring in spatiotopic regions, whereas the other (LC) may be essentially non-spatial and more related to temporal relations between events and the recruitment of resources (effort or arousal). While coherent with the roles associated to these neuromodulatory systems, the ventral attentional network (which we found to be involved during spatial interference), has been proposed to recruit LC after detection of salient events (Corbetta et al., 2008); however, since we did not observe that LC was related to changes in spatial interference, its role may mainly relate to non-spatial aspects of the attention network.

### 4.5 Individual differences

The analysis of individual differences showed that, across studies, load and crowding exerted different effects on participants’ accuracy. In low crowding, low performers had a larger decrease in accuracy than high performers in both studies. The effect of crowding was significant in the pupillometry but not in the fMRI study. This result suggests that the processes of load and crowding are separate in terms of compromising participants’ performance, something that has been already proposed (Intriligator & Cavanagh, 2001; Oksama & Hyönä, 2004). The pupil results showed that high performers showed pupil dilation with load both in low and high crowding, suggesting that they were able to allocate resources even in the most difficult condition. The brainstem analysis on individual differences indicated that, in addition to being recruited with increasing load, the LC was also to some extent recruited with increasing crowding but only in high performing participants. Together, these results seem to reflect these participants’ ability to flexibly recruit resources to keep high levels of accuracy in a highly challenging condition. Thus, our findings suggest the intriguing notion that high performers are not only better prepared to resolving high conflict (i.e. high load) trials, but also able to recruit most effectively their cognitive resources during cognitive conflict.

Together with the cholinergic systems, the dopaminergic and noradrenergic systems modulate and facilitate visuo-spatial attention and visual working memory processes (Sarter et al., 2006; Störmer et al., 2012). Individual differences are related to genetic and physiological variations in these systems, and they may underlie variations in aging as well as states of mental illnesses (Mather & Harley, 2016; Störmer et al., 2012; Unsworth & Robison, 2017).

### 4.5 Relation with theories on limitations in dynamic visual attention

Theoretically, two competing models have been proposed to account for the perceptual and cognitive limitations observed during a task requiring dynamic attention, such as MOT (Alvarez & Cavanagh, 2004; S. L. Franconeri et al., 2013). One is the coding of individual stimuli into cortical spatial maps that may inhibit each other due to close perceptual proximity of the stimuli (cortical real-estate account). Another is the coding of stimuli in separate working memory slots (“resource” account). While the first predicts that load and crowding engage the same substrates in primary visual circuits, the second would predict that different networks deal with load and crowding (that is, objects are represented as informational chunks and additional mechanisms that deal with spatial interference due to crowding are recruited as needed). Although our results do support the cortical real-state account, given that spatial interference influences accuracy and cortical as well as some of the brainstem activations, this view encounters difficulty in explaining effects on accuracy, cortical and brainstem activation, and pupil dilations when spatial interference is held constant. Moreover, we observed activations in LC and pupil responses related to load and not to the spatial aspects of crowding. Hence, a general “resource” account as originally proposed by Kahneman (Kahneman, 1973), although it may be unsatisfactory for a number of reasons (S. L. Franconeri et al., 2013), remains a candidate explanation. This agrees with empirical results using object-tracking with circular trajectories and varying degrees of separation (Holcombe et al., 2014). However, we note that the above two views have mainly focused on the perceptual level. Previous studies suggest that not only perceptual but also executive processing is crucial for resolving MOT, not only during tracking but also at the moment of target selection (Allen et al., 2006; Eayrs & Lavie, 2018; S. L. Franconeri et al., 2007; Lapierre et al., 2017; Ma & Flombaum, 2013; Tombu & Seiffert, 2008; Vul et al., 2009). Therefore, a resolution between the two hypotheses is not straightforward based on the present results. Nevertheless, MOT shares many characteristics with other visuo-spatial tasks that rely on perceptual as well as executive processes (i.e. VSTM, (Fougnie & Marois, 2006), (Marois & Ivanoff, 2005); subitizing: (Chesney & Haladjian, 2011); selection of objects: (Xu & Chun, 2009)) and consequently, its study may inform how limitations in the other processes occur.

### 4.6 Study limitations

For the present work, we only had two levels of crowding (i.e. spatial interference). Future experiments with more levels of crowding in addition to load would allow a more complete picture of the relation between the two factors. Although we found no interaction between them, adding crowding levels may present a more complex picture. Another remaining question is what would be the effect of informing participants at the start of the trial whether there will be high or low crowding in the display at the same time as telling the number of targets. A prediction would be that, if preparatory activity related to load would be engaged to prepare for higher crowding, the accuracy patterns of load and crowding would be dependent (Figure 1.c). For the purpose of this study, we focused on the role of attention in resolving spatial interference during tracking and therefore did not study brain and brainstem activity during visual selection (i.e. activity during the target assignment phase). Studies focusing on this stage may confirm previous results suggesting that executive functions are involved in this stage and influence tracking, and extend our results regarding the involvement of brain and brainstem regions in the encoding of targets.

## 5. Conclusion

Our results support the view that, in the MOT task, distinct mechanisms deal with the increasing demands posed by cognitive workload (i.e. number of target objects) and spatial interference (i.e. proximity between targets and distractors), and brainstem neuromodulatory nuclei play a key role. Load, additionally to spatial interference, compromised accuracy during MOT. A set of regions within the dorsal attention network were recruited with both load and spatial interference demands. Additional areas within the ventral attention network were recruited during high crowding. The ventral and dorsal attentional systems collaborate to solve the challenges of covert goals, likely engaging top-down control, as well as rising demands, likely engaging bottom-up processes (Corbetta et al., 2008; Vossel et al., 2014). Locus coeruleus and pupil dilation signaled effort in the sense of engagement, which occurred with increasing load and, in high performers, with higher crowding. VTA/SN differed from LC in that it was related to spatial interference. The MOT task, together with physiological measures, may offer a unique opportunity to interrogate how cortical and neuromodulatory systems are adaptively recruited when cognition is challenged.

